# Customizable bead-based platform for EV surface protein immunocapture and detection in plasma

**DOI:** 10.64898/2026.10.28.762484

**Authors:** Hema Saranya Ilamathi, Mariana Castañeda Mesa, Metoboroghene O. Mowoe, Srinidhi Rangarajan, Doste Rashid Mamand, Oliver George Hayes, Mattias Hällbrink, Malgorzata Honcharenko, André Görgens, Daniel W. Hagey, Oscar P.B Wiklander

## Abstract

Extracellular vesicles (EVs) are increasingly recognized as valuable biomarkers for liquid biopsy because they carry molecular signatures that reflect their cells of origin. However, EV isolation remains a challenge, and existing EV immunocapture platforms are constrained by nonspecific binding, limited flexibility, and insufficient sensitivity. Here, we present a customizable bead-based platform for the isolation and detection of specific EV populations directly from plasma. The platform combines bead-immobilized capture antibodies with fluorescent detection antibodies, enabling sensitive and multiplexed EV profiling. Optimization of bead conjugation chemistry improved antibody immobilization and reduced nonspecific background signals, enhancing assay specificity and performance. Using breast cancer-derived EVs expressing fluorescent protein, we demonstrate the detection of tumor-derived EVs in plasma at concentrations as low as 1 × 10^9^ EVs/ml. Furthermore, the assay distinguished HER2- and TROP2-high from low EV populations, demonstrating its ability to detect clinically relevant differences in biomarker expression. The workflow is compatible with direct EV analysis in the unprocessed plasma samples, thereby reducing sample preparation requirements and assay complexity. These findings establish a sensitive, flexible, and scalable platform for EV biomarker analysis. The approach overcomes several limitations of current commercial bead-based technologies and may facilitate the development of customizable multiplexed EV-based assays for cancer detection, patient stratification, and disease monitoring.

## Introduction

Liquid biopsy has emerged as a promising tool for disease diagnosis, prognosis, and therapeutic monitoring, as it enables minimally invasive sampling and repeated longitudinal assessment of patients over time ^1–3^. Tissue biopsies provide important molecular information but are invasive, prone to sampling bias, and difficult to repeat during treatment monitoring ^1–3^. Biological fluids such as blood contain a diverse range of circulating biomolecules, including nucleic acids, proteins, lipids, metabolites, and extracellular vesicles (EVs), many of which reflect the physiological and pathological status of their tissue of origin and offer the possibility of tracking the disease status in a minimally invasive way^1–3^ . Among the analytes, circulating tumor DNA (ctDNA) has recently been introduced into clinical routine to monitor the disease. However, ctDNA is often present at low abundance in early-stage disease ^4^. EVs have attracted considerable interest as biomarkers to overcome the limitations of ctDNA, as they protect their molecular cargo from degradation and carry proteins, nucleic acids, lipids, and metabolites that mirror the characteristics of the cells from which they are released ^5, 6^. Alterations in EV abundance and cargo composition have been associated with a wide range of diseases, including cancer, neurodegenerative disorders, cardiovascular diseases, and inflammatory conditions ^6–10^. In cancer, tumor-derived EVs play important roles in intercellular communication and contain clinically relevant biomarkers that may support early detection, patient stratification, and treatment monitoring ^6, 11^.

Despite their diagnostic potential, the reliable isolation of EVs from complex biological fluids remains challenging. Conventional EV isolation methods, including differential ultracentrifugation, density-gradient ultracentrifugation, size-exclusion chromatography (SEC), tangential flow filtration, and membrane filtration, are widely used in research settings ^12–14^. However, these approaches often involve lengthy sample processing workflows that are time consuming, expensive, require specialized instrumentation, and introduce variability in EV yield and purity ^12, 13, 15^. Furthermore, all these methods co-isolate soluble proteins, lipoproteins, protein aggregates, and other extracellular particles, which can interfere with downstream analyses and reduce assay specificity^15,16^. Although density-gradient centrifugation can yield highly purified EV preparations, its complexity and low throughput make it unsuitable for routine clinical applications ^17^. In contrast, polymer-based precipitation methods offer operational simplicity but extensively coprecipitate non-vesicular components, compromising analytical accuracy.

To overcome these limitations, affinity-based approaches have been developed to selectively isolate EVs based on the expression of specific surface-associated proteins ^18, 19^. Antibody-mediated capture strategies enable the enrichment of specific EV subpopulations and facilitate the analysis of disease-associated biomarkers directly from biological samples ^20–28^. Several commercial platforms have adopted this principle for EV profiling; however, important challenges remain. The commercial bead-based assays rely on predefined antibody panels, limiting flexibility in biomarker selection and restricting adaptation to newly identified or disease-specific targets ^26, 28^. These constraints are particularly relevant for cancer diagnostics, where biomarker composition can vary substantially between tumor types, molecular subtypes, and disease stages ^29–31^. In addition, detecting less abundant disease-associated EV populations directly in complex biological fluids such as plasma remains difficult because of background particles, soluble proteins, and limited assay sensitivity. Furthermore, EV characterization platforms such as nanoparticle flow cytometry (NanoFCM) and ExoView require specialized instrumentation and are associated with higher per-sample costs than conventional flow cytometry-based assays, which may limit their widespread clinical implementation. Consequently, there remains a need for bead platforms that combine high specificity, customizable capture-detection strategies, direct compatibility with unprocessed biofluids, and implementation on standard flow cytometers. Such systems could simplify EV biomarker analysis while improving adaptability to different disease settings and facilitating clinical translation.

Here, we present a customizable bead-based platform for the sensitive and specific detection of EV-associated protein biomarkers. By optimizing bead functionalization chemistry and implementing a dual-antibody detection strategy, we achieve efficient EV capture with minimal background binding. The platform enables flexible selection of capture and detection antibodies, supporting multiplexed biomarker analysis. Using engineered breast cancer-derived EVs and plasma spike-in model experiments, we demonstrate the direct detection of tumor-derived EVs in unprocessed plasma and the discrimination of clinically relevant HER2 expression levels. Collectively, our results establish a rapid, scalable, and adaptable approach for EV biomarker profiling that addresses key limitations of existing bead-based technologies and may facilitate the development of clinically applicable EV-based liquid biopsy assays.

## Methods Cell culture

SKBR3, MDA MB 231, T47D, and CD81mNG-overexpressing engineered SKBR3 cell lines were grown in Dulbecco’s Modified Eagle Medium (DMEM) supplemented with 10% fetal bovine serum and 1% Penicillin-streptomycin. Cultures were incubated at 37 °C with 5 % CO_2_. All cell lines were regularly monitored for mycoplasma.

## Human blood samples

Anonymous blood samples from healthy donors were purchased from Karolinska University Hospital. All samples were received after signed informed consent. The study was performed in accordance with the Declaration of Helsinki and approved by the Ethical Review Board in Stockholm (2017/912-31) and the Swedish Ethics Board (2024-00754-02).

## EV production and preparation/enrichment

10 x 10^6^/ml cells were seeded into 150 mm dishes and expanded in DMEM until they reached 70-80% confluency. Cells were washed with PBS before switching to OPTIMEM media supplemented with 1% Penicillin-streptomycin. After 48 hours, conditioned media were collected and centrifuged at 700 g for 5 min and 2000 g for 10 min to remove cells and debris. The supernatant was filtered further through a 0.22- micron filter. The conditioned media were concentrated to 1 mL using 10 kDa ultra-centrifugal filters and centrifuged at 4000g for 30 minutes each round and maintained at 4°C. Concentrated conditioned media were then processed with size exclusion chromatography (qEV Izon 70 nm) for EV isolation, removing proteins and further contaminants. Columns were eluted sequentially with 1 mL of PBS, and fractions 4 and 5 were collected for further concentration to 1 mL with 10 kDa filters. EVs were stored in HAT buffer at –80 °C for later use.

## Nanoparticle tracking analysis

EV particle concentration and size distribution were determined using a Zetaview instrument (Particle Metrix GmbH). Samples were diluted in PBS (Gibco) to a suitable concentration for nanoparticle tracking analysis and were analyzed in 11 different fields at 25 °C with a camera sensitivity setting at 80 and a threshold at 100. Recordings were analyzed using Zetaview software 8.05.16 SP3. Particle counts were corrected for measurements of PBS, followed by correction for dilution.

## Carbodiimide-based bead platform

PMMA (#AR146, Microparticles GmbH) beads (1:4000) were initially washed in PBST buffer (PBS with 0.05% Tween 20). To minimize bead loss caused by adherence to the tube surface, all experiments were performed using polystyrene tubes (Fisher Scientific, #10186400). Beads were pelleted at 2000g for 5 minutes at room temperature (RT) and resuspended in PBST. Next, beads were activated with 0.3 M EDAC (1-Ethyl-3-(3-dimethylaminopropyl) carbodiimide) in 100mM MES buffer, pH 6.0 (2-(N-morpholino) ethanesulfonic acid). The buffer and EDAC concentration were optimized to efficiently immobilize capture antibodies on beads (**Figure S1A & B**). The beads were kept in a shaker for 20 minutes at RT. The beads were pelleted and resuspended in PBS. The capture antibody was added as specified in the experiments and incubated overnight at 4℃ in a shaker. The reaction was stopped with Borate buffer, pH 8.0 (100 mM boric acid, 75 mM sodium chloride, 25 mM sodium tetraborate), and the beads were pelleted. Next, the beads were blocked in PBS with 3% Bovine Serum Albumin (BSA) for 2 hours at RT on a shaker. Beads were spun down and resuspended in PBS before adding the desired concentration of EVs, as mentioned in the experiments. The beads were incubated with EVs at 4℃ in a shaker for 2 hours. The incubation time was optimized to efficiently capture EVs on beads (**Figure S1C**). Then, EV-bound beads were pelleted, and a fluorescent-tagged detection antibody was added. After incubating with the detection antibody for 10 minutes in a shaker at RT, beads were pelleted and resuspended in PBS. The beads were analyzed by the MacsQuant analyzer 16. The flow rate was maintained low with sample mixing before analysis, and 25 µL of sample was analyzed per condition.

## Flow cytometry

The EVs captured on beads were measured by the MACS Quant analyzer 16. The following monoclonal antibodies were used for either capturing or detecting EVs: CD63 PE (Miltenyi Biotec, #130-118-077, clone REA1055), CD81 PE (Miltenyi Biotec, #130-118-342, Clone REA513), CD9 PE (Miltenyi Biotec, #130- 123-761, Clone SN4 C3-3A2), CD63 APC (Miltenyi Biotec, #130-118-078, Clone REA1055), CD81 APC (Miltenyi Biotec, #130-119-787, Clone REA513), Trastuzumab (clinical grade from pharmacy), Sacituzumab (#HY-P99045, MedChem Express). Flow cytometric data were analyzed using the FlowJo software version 10.10.0 (FlowJo LLC). Fluorescence data were transformed using a scale to accommodate low and negative values while preserving the log-like behavior of the data at higher intensities. Under these conditions, geometric mean fluorescence intensity (gMFI) was used to summarize signal, as it provides a representative measure of central tendency for log-distributed data and is sensitive to shifts in fluorescence intensity across conditions. Median-based analysis also produced consistent trends (data not shown).

## EV spike-in plasma

Different concentrations of EVs were spiked into plasma (1:1 diluted in PBS) and added directly to PMMA beads. For HER2 and TROP2 biomarker detection, EVs at a concentration of 1E10/ml were spiked into the plasma. The beads were incubated with spiked-in plasma or isolated EVs for 2 hours, and the rest of the procedure was carried out as mentioned above in the protocol.

## Statistical analysis

The graphs were plotted, and statistical analysis was carried out using GraphPad Prism 10.6.1. Depending on the experiment, conditions were compared by two-way or one-way ANOVA with Tukey’s, Dunnett’s, or Šidák’s multiple comparison test. The statistically significant population was highlighted in the graph.

## Results

### 1. Development and optimization of bead-based EV capture platforms

To enable sensitive detection of EV-associated surface proteins, we developed a bead-based platform that combines antibody-mediated EV capture with flow cytometric analysis (**Fig. 1**). The assay relies on immobilization of a capture antibody on the bead surface, followed by incubation with EV-containing samples and detection using a fluorescently labelled antibody directed against a second EV surface marker. This dual-antibody strategy enables selective EV detection and flexible biomarker profiling.

**Figure 1.**
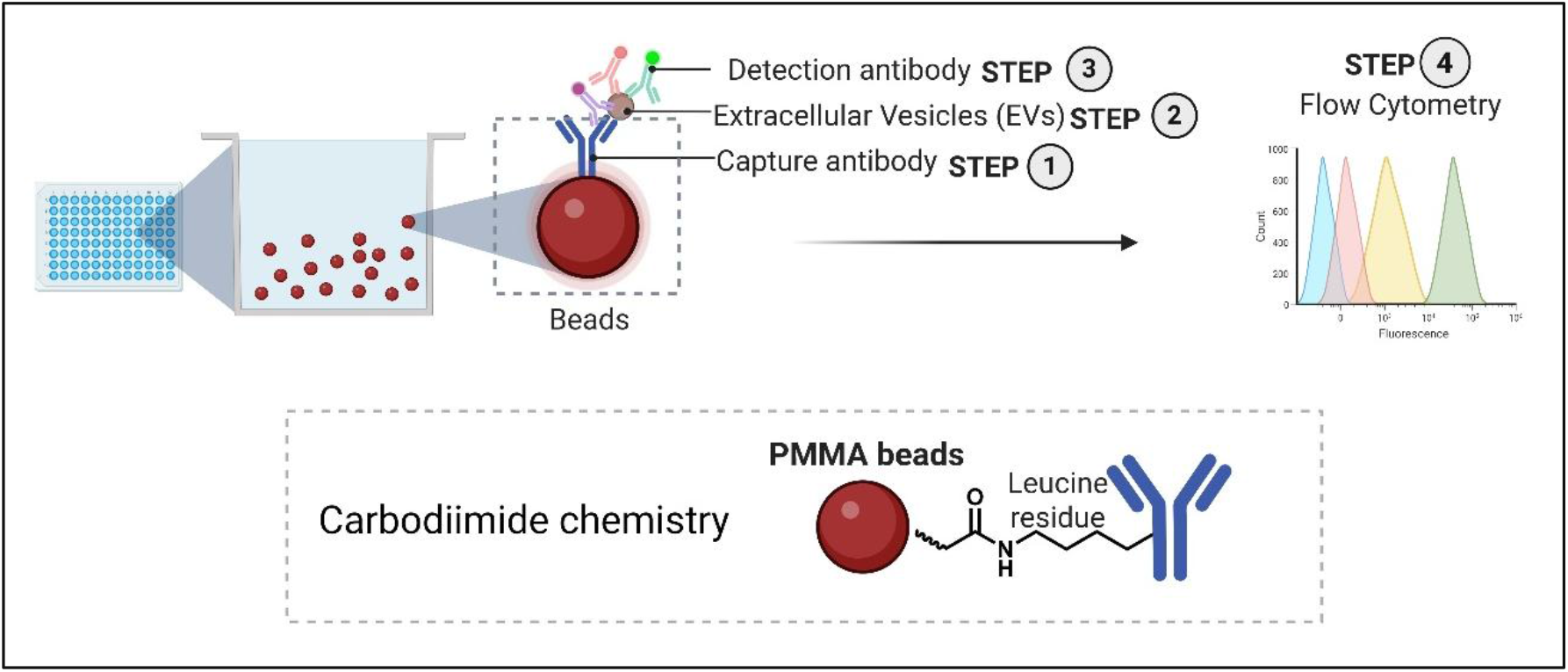
Schematic representation of the bead platform. Overview of the bead-based EV assay workflow, including the chemistry used for antibody conjugation to the bead surface.

**Supplementary Figure 1.**
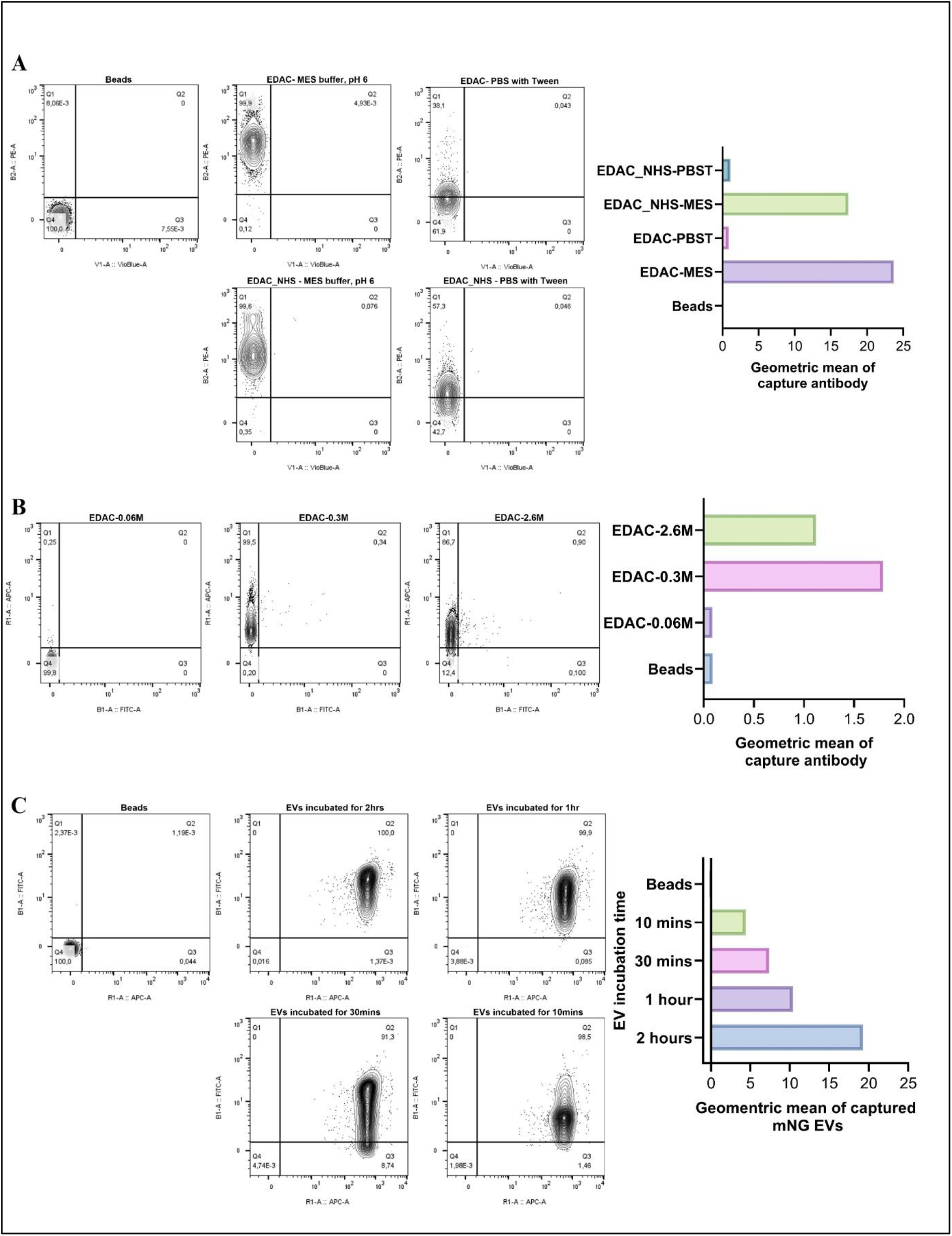
: Optimization of PMMA bead functionalization for antibody immobilization. A) Optimization of PMMA bead activation conditions. Beads were activated using either EDAC alone or EDAC in combination with NHS in MES buffer (pH 6.0) or PBS containing Tween-20 (pH 7.0), followed by antibody conjugation. B) Effect of EDAC concentration on PMMA bead activation in MES buffer (pH 6.0). Graphs show gMFI of immobilized capture antibodies detected using CD63-PE or CD9-APC. C) SKBR3 CD81 mNG EVs were incubated for different times (10 minutes, 30 minutes, 1 hour, 2 hours) with CD81 APC-coated PMMA beads. The bar plot shows the gMFI of the captured mNG EVs. Data are representative of a single experiment.

Poly(methyl methacrylate) (PMMA) beads functionalized with carboxyl groups were conjugated to capture antibodies using EDAC-mediated carbodiimide chemistry (**Fig. 1**). Conditions for bead activation, antibody coupling, and EV capture were systematically optimized (**Supplementary Fig. S1**). Bead activation was first evaluated in different buffers by assessing capture-antibody binding efficiency. Activation in MES buffer (pH 6.0) resulted in more efficient and homogeneous antibody immobilization than activation in MES buffer containing NHS (N-hydroxysuccinimide), whereas activation in PBS containing Tween-20 (pH 7.0) produced minimal antibody binding, indicating poor bead activation (**Fig. S1A**).

We next optimized the EDAC concentration used for bead activation. Antibody immobilization efficiency was dependent on EDAC concentration, with optimal coupling achieved at 0.3 M. Lower or higher EDAC concentrations resulted in reduced antibody binding (**Fig. S1B**). Finally, incubation times for EV capture were evaluated using capture antibody-coated beads. EV capture efficiency increased with incubation time and reached a maximum after 2 h, whereas shorter incubation periods resulted in lower EV binding (**Fig. S1C**). These optimized conditions were used throughout the subsequent bead-based EV capture workflow.

### 2. Specificity of the platform

To validate assay specificity, we first confirmed successful functionalization of PMMA beads with capture antibodies and assessed potential background arising from non-specific or cross-reactive binding of the detection antibody. Detection antibody signal was observed only when EVs were present, indicating specific target recognition (**Fig. 2A**). We then examined non-specific interactions between EVs and the PMMA bead surface. In the absence of capture antibodies, 3% BSA blocking effectively prevented non-specific EV adsorption and reduced background signal to baseline levels (**Fig. 2B**).

**Figure 2.**
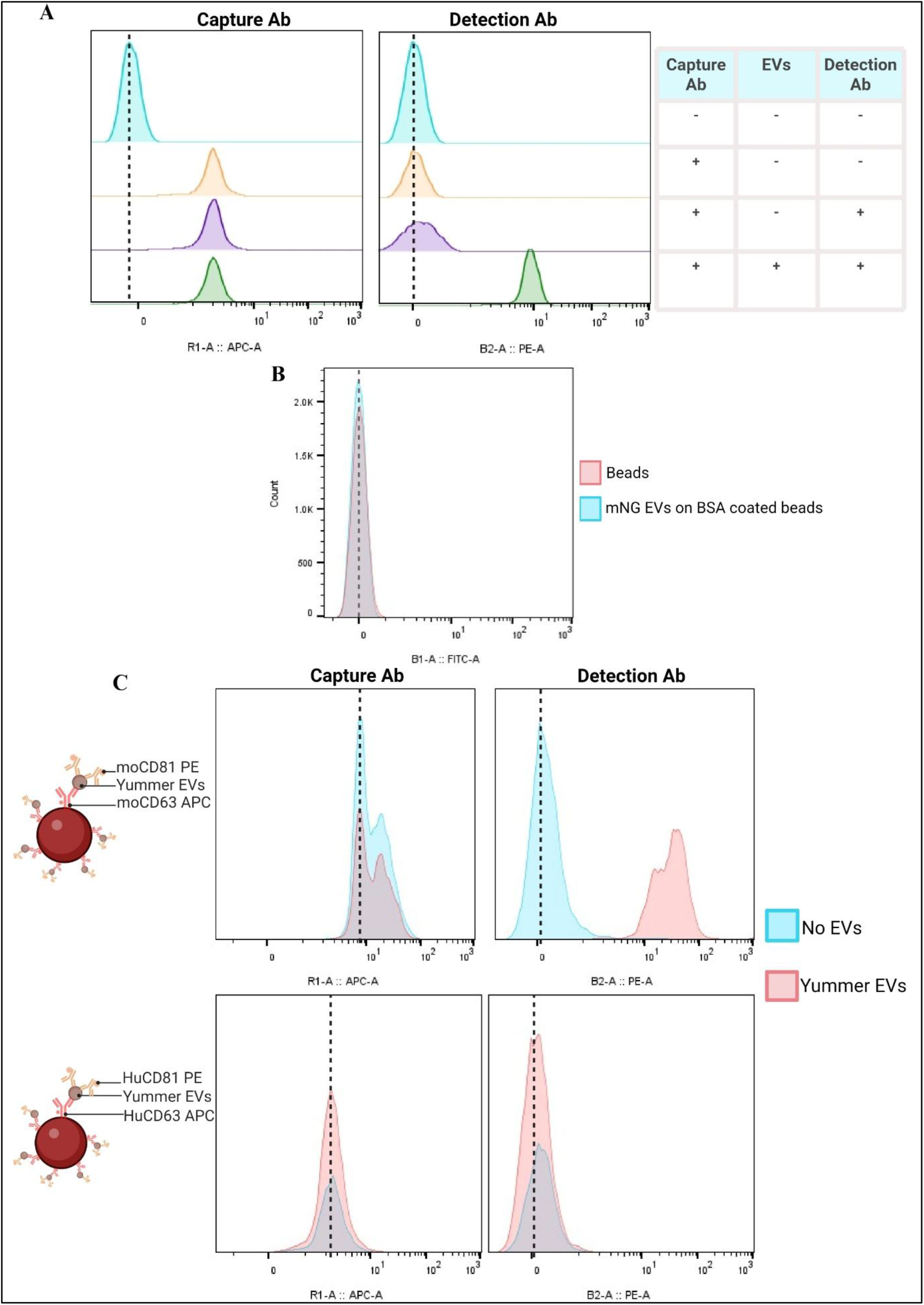
Specificity of the bead-based EV capture platform. (A) Assessment of nonspecific binding using anti-CD63 APC-functionalized PMMA beads and anti-CD81 PE detection antibody in the presence or absence of SKBR3-derived EVs. Representative histograms from three independent experiments. (B) Blocking with 3% BSA reduces nonspecific EV binding and background signal. (C) Species-specific validation using EVs from mouse Yummer cells. EVs were captured and detected with mouse- or human-specific antibody pairs, demonstrating selective detection only with species-matched antibodies. Representative histograms from three independent experiments.

We next assessed the specificity of antibody-mediated EV capture using EVs derived from the mouse melanoma cell line, Yummer. Beads coated with human-specific CD63 or CD81 antibodies failed to generate detectable signals when incubated with mouse-derived EVs and subsequently stained with human-specific detection antibodies (**Fig. 2C**). These findings demonstrate that EV detection was dependent on specific antibody-antigen interactions rather than nonspecific binding events. Collectively, these data confirm that both platforms selectively capture target EV populations while maintaining low background signal.

### 3. Sensitivity of the system

To optimize EV capture, we systematically evaluated the impact of bead concentration and capture antibody density using engineered SKBR3-derived CD81-mNeonGreen (mNG) EVs as a fluorescent model system (**Fig. 3A-C**). Quantification of the mNG signal showed that both parameters strongly affected capture performance, enabling identification of optimal assay conditions (**Fig. 3B-C**). PMMA beads diluted 1:4,000 and functionalized with 0.5 μg mL⁻¹ capture antibody were selected for further analyses based on EV capture efficiency (**Fig. 3C**). Using these conditions, we assessed assay sensitivity by measuring EV detection across a range of input concentrations. The platform reliably detected EVs at concentrations as low as 5 × 10⁸ EVs mL⁻¹ (**Fig. 3D-F**). Together, these results highlight the sensitivity of the PMMA bead-based platform for EV detection and support its suitability for downstream biomarker analyses.

**Figure 3.**
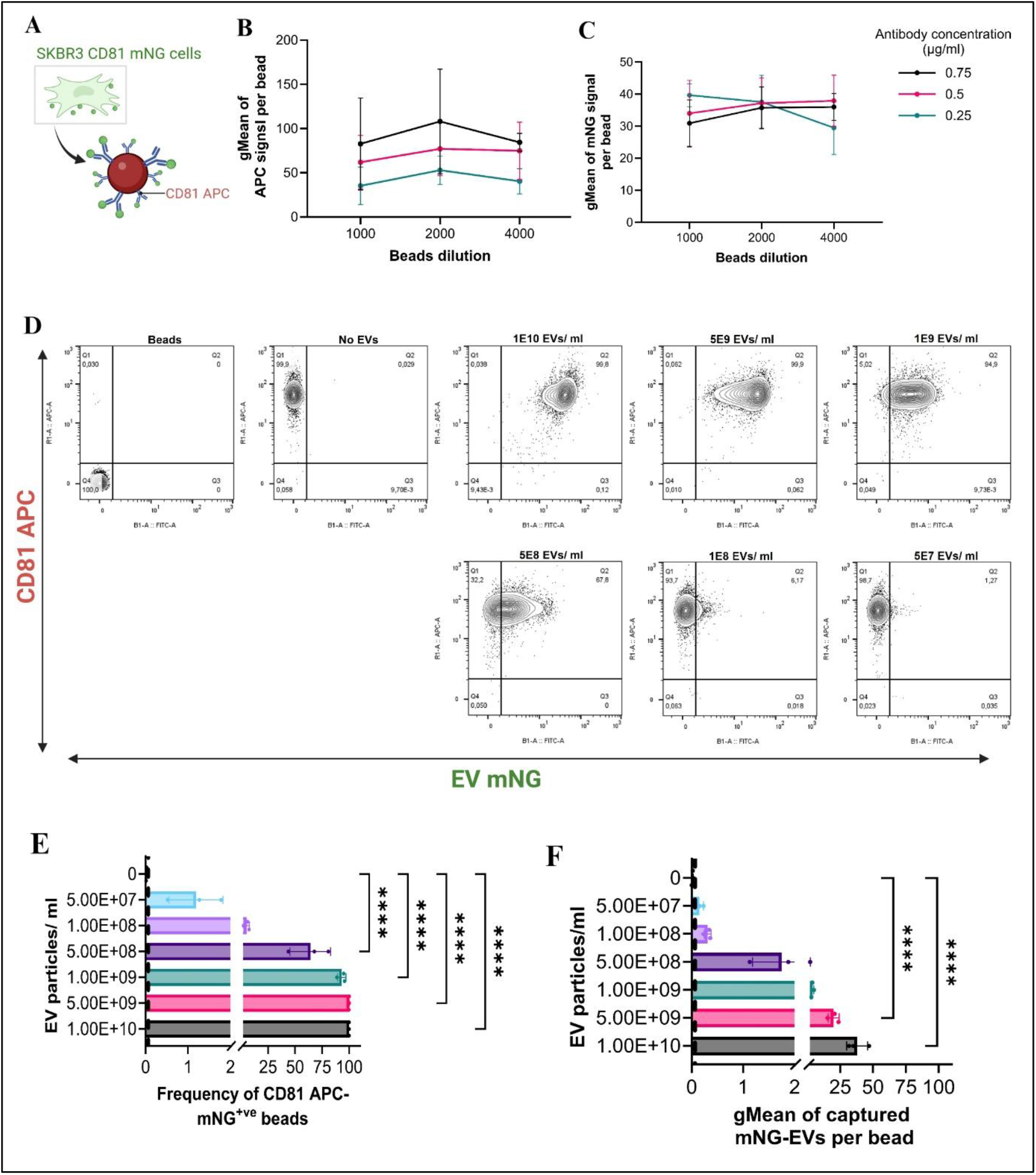
Optimization and sensitivity of the bead-based EV capture platform. (A–C) PMMA beads coated with increasing concentrations of anti-CD81 capture antibody and tested at different bead dilutions were incubated with SKBR3-derived mNeonGreen-positive (mNG⁺) EVs (1 × 10¹⁰ EVs mL⁻¹). Representative plots (A), capture antibody density per bead (gMFI; B), and EV capture signal (mNG gMFI; C) are shown. No significant differences were observed (two-way ANOVA with Tukey’s multiple-comparisons test). (D–F) Sensitivity assessment using PMMA beads diluted 1:4,000 and coated with 0.5 μg mL⁻¹ anti-CD81 antibody. SKBR3-derived mNG⁺ EVs were analyzed across a concentration range of 1 × 10¹⁰ to 5 × 10⁷ EVs mL⁻¹. Representative contour plots (D), frequency of mNG⁺ beads (E), and mNG gMFI per bead (F) are shown. Data were analyzed by one-way ANOVA with Tukey’s multiple-comparisons test. Data are representative of three independent experiments and presented as mean ± s.d. ****P < 0.0001, ***P < 0.001, **P < 0.01, *P < 0.05.

### 4. Direct detection of plasma-derived and tumor-derived EVs

Having established robust EV detection using cell culture-derived EVs, we next evaluated whether the PMMA bead platform could detect EVs directly in human plasma without prior EV isolation. PMMA beads functionalized with anti-CD81 capture antibodies efficiently enriched plasma-derived EVs, as confirmed by detection with anti-CD63 antibodies (**Fig. 4A**). In contrast, beads coated with anti-CD63 capture antibodies produced substantially lower signals, indicating lower CD63-positive EVs in the healthy plasma sample (**Fig. 4A**).

**Figure 4.**
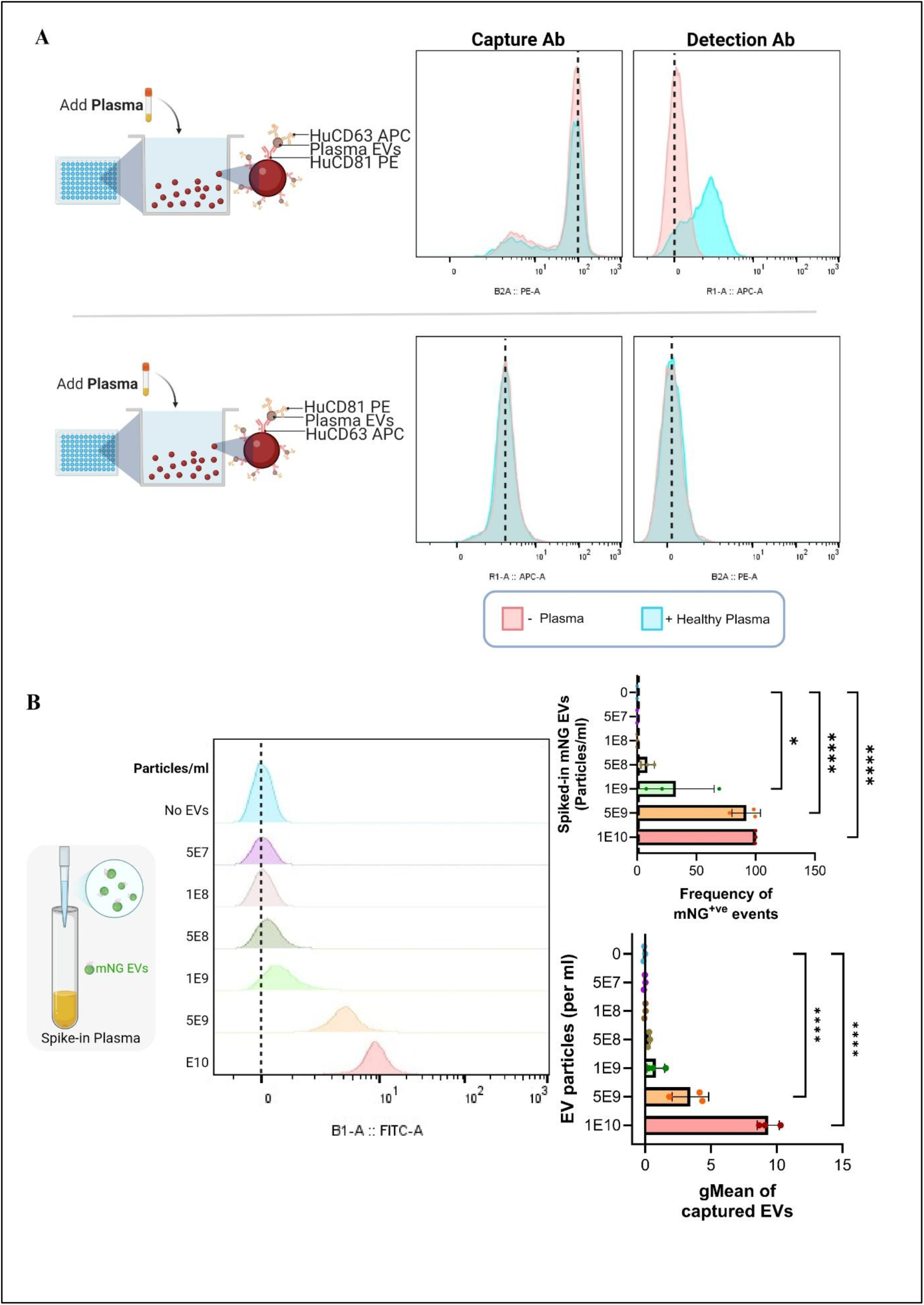
Direct detection of plasma-derived EVs from unprocessed plasma. (A) EVs were captured directly from unprocessed plasma using PMMA beads coated with anti-CD63 or anti-CD81 antibodies and detected with the reciprocal antibody. EV detection was enhanced when anti- CD81 was used for capture. (B) Detection of tumour-derived EVs in plasma was evaluated by spiking engineered SKBR3-derived CD81 mNeonGreen-positive (mNG⁺) EVs at increasing concentrations. Representative histograms show mNG⁺ EVs captured on CD81-decorated PMMA beads. Graphs display the percentage of mNG⁺ EV-positive beads and the mNG gMFI per bead. Data are representative of three independent experiments and are presented as mean ± s.d. Statistical analysis was performed using one-way ANOVA with Dunnett’s multiple-comparisons test. ****P < 0.0001, **P < 0.01, *P < 0.05.

We then assessed the ability of the platform to detect tumor-derived EVs in a complex biological matrix. Engineered SKBR3-derived CD81-mNeonGreen (mNG) EVs were spiked into human plasma at varying concentrations and analyzed directly using the PMMA bead-based assay. Tumor-derived EVs were readily detected across the tested concentration range, with reliable detection observed down to 1 × 10^9^ EVs mL⁻¹ (**Fig. 4B**). Both the frequency of mNG-positive beads and the fluorescence intensity per bead decreased in a concentration-dependent manner, consistent with reduced EV input (**Fig. 4B**).

Collectively, these findings demonstrate that the PMMA bead platform enables sensitive detection of both endogenous plasma EVs and tumor-derived EVs directly from unprocessed plasma, without the need for prior EV isolation or enrichment.

### 5. Detection of clinically relevant EV-associated biomarkers in plasma

To determine whether the PMMA platform could distinguish EV populations based on clinically relevant biomarkers, we analysed HER2 and TROP2 expression on breast cancer-derived EVs. PMMA beads were conjugated to clinical antibodies such as Trastuzumab or Sacituzumab that target HER2 and TROP2, respectively (**Figure 5A, B**). For HER2 detection, PMMA beads were functionalized with trastuzumab and incubated with plasma samples spiked with EVs derived from HER2^high^ SKBR3 cells or HER2^low^ MDA-MB-231 cells (**Fig. 5A**). Trastuzumab-conjugated beads could distinguish EVs derived from SKBR3 and MDA-MB-231 cells that are HER2^high^ and HER2^Low^, respectively (**Figure 5A**). We next evaluated TROP2 detection using PMMA beads functionalized with sacituzumab. Plasma samples spiked with EVs derived from SKBR3 cells and T47D cells were analysed under identical conditions (**Fig. 5B**). SKBR3-derived EVs exhibit higher TROP2 expression than T47D-derived EVs (**Figure 5B**). Overall, these findings demonstrate that the PMMA bead platform supports direct detection of clinically relevant EV-associated biomarkers in plasma and enables discrimination between EV populations with distinct molecular phenotypes.

**Figure 5.**
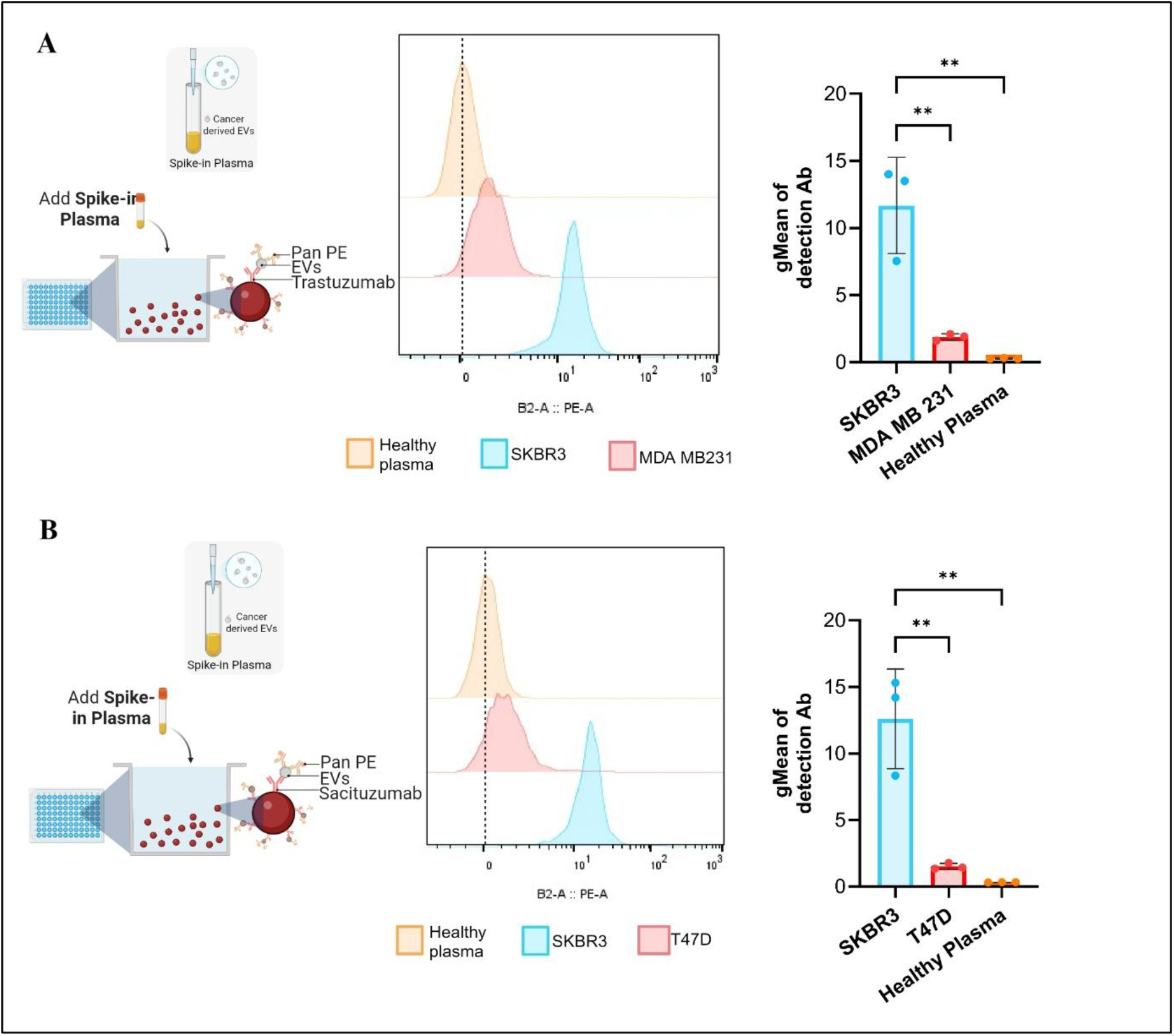
Detection of tumor-associated EV biomarkers in plasma. (A) HER2-positive EVs were detected in plasma spiked with HER2^high^ SKBR3-derived or HER2^low^ MDA-MB-231-derived EVs (1 × 10¹⁰ EVs mL⁻¹). EVs were captured on trastuzumab-coated PMMA beads and detected using Pan PE (CD63, CD81, CD9) antibodies. Representative histograms and EV-associated fluorescence intensity (gMFI) per bead are shown. (B) TROP2-positive EVs were detected in plasma spiked with SKBR3- or T47D-derived EVs (1 × 10¹⁰ EVs mL⁻¹). EVs were captured on sacituzumab-coated PMMA beads and detected using Pan PE (CD63, CD81, CD9) antibodies. Representative histograms and EV-associated fluorescence intensity (gMFI) per bead are shown. Data are representative of three independent experiments and presented as mean ± s.d. Statistical analysis was performed using one-way ANOVA with Šidák’s multiple-comparisons test. P < 0.01.

## Discussion

Bead-based approaches have become widely used for extracellular vesicle (EV) characterization because they overcome the detection limitations imposed by the small size of EVs and enable analysis using conventional flow cytometers. Current platforms range from immunocapture-based magnetic beads for EV enrichment to multiplexed systems such as MACSPlex, which facilitate broad profiling of EV surface markers. These technologies have significantly expanded EV phenotyping capabilities but often involve trade-offs between assay flexibility, cost, sensitivity, multiplexing capacity, and compatibility with complex biological samples^26, 32, 33^.

Unlike multiplexed bead arrays that rely on predefined antibody panels, the platform allows independent selection of capture and detection antibodies, enabling selective enrichment of specific EV populations while simultaneously interrogating additional surface biomarkers. This modular design provides flexibility for biomarker discovery and validation studies, where the relevant EV-associated markers may differ between diseases and are often unknown at the beginning of an investigation. In contrast to fixed-panel approaches, the assay can be readily adapted by changing antibody combinations without altering the underlying workflow.

A second important distinction is the ability to perform EV analysis directly in unprocessed plasma. Many bead-based EV assays achieve robust performance only after EV enrichment through ultracentrifugation, size-exclusion chromatography, precipitation, or affinity purification^32, 33^. While these approaches improve EV purity, they increase workflow complexity, require higher volumes of samples, and may introduce sample loss or selection bias. In the present study, PMMA beads efficiently captured endogenous plasma EVs and detected tumour-derived EVs spiked into plasma without prior isolation. This suggests that the platform may preserve EV heterogeneity while reducing sample handling requirements, which is advantageous for both large-scale biomarker studies and future clinical implementation.

The analytical performance of bead-based EV assays is strongly influenced by biomarker abundance. Previous studies using multiplexed bead platforms have reported detection of highly abundant tetraspanins at relatively low EV concentrations, whereas detection of less abundant surface proteins typically requires substantially higher EV input^26^. Consistent with these observations, sensitivity in our system depended on the biomarker used for detection. Importantly, assay sensitivity in this study was evaluated using engineered mNeonGreen-positive EVs, where signal intensity depends not only on EV concentration but also on the proportion of EVs carrying detectable fluorescent cargo. Consequently, the reported detection limit likely represents a conservative estimate of platform performance. Future studies evaluating endogenous EV markers and clinical samples will provide a more comprehensive assessment of analytical sensitivity across different biological contexts.

Compared with existing bead-based approaches that primarily quantify canonical EV markers, the PMMA platform enables selective interrogation of disease-associated EV subpopulations. We show here that HER2 is released in EVs in a manner that reflects the parental cellular HER2 expression profile. In contrast, TROP2 displays a different pattern, despite T47D cells being known to express higher levels of TROP2. This observation suggests that EV protein cargo may not always directly mirror cellular protein abundance and that EV protein packaging may vary depending on both the cell type and the biomarker examined^34–36^. Further studies are needed to better understand the biological mechanisms underlying these differences. Overall, the ability to distinguish clinically relevant biomarkers on EV populations in plasma is particularly relevant for liquid biopsy applications, where tumour-derived EVs typically represent only a small fraction of the total circulating vesicle pool. The possibility of combining target-specific capture with downstream phenotyping may therefore provide greater biological and clinical resolution than approaches focused solely on bulk EV measurements^32^.

A further practical advantage of the platform is its compatibility with conventional flow cytometers. Single-EV analysis technologies, including nanoscale flow cytometry and specialized optical platforms, can provide higher analytical resolution but often require dedicated instrumentation and technical expertise. By leveraging standard flow cytometry infrastructure, the PMMA platform offers a more accessible route for EV biomarker analysis while maintaining sufficient flexibility to accommodate different biomarker combinations and experimental questions.

In summary, the PMMA bead platform combines several features that are not typically integrated within a single bead-based EV assay, including customizable capture-detection strategies, direct compatibility with unprocessed plasma, and implementation using conventional flow cytometry. By bridging selective EV enrichment and flexible biomarker interrogation, the platform addresses key limitations of current bead-based technologies and provides a practical framework for EV biomarker discovery, patient stratification, and liquid biopsy development.

### Limitations of the study

This study demonstrates that a PMMA-based platform can be customized for the direct analysis of extracellular vesicles (EVs) in plasma. However, this study has some limitations. First, platform performance was evaluated primarily using engineered EVs and plasma spike-in models. Although these systems provide controlled conditions for proof-of-concept validation, studies involving larger and more diverse patient cohorts are necessary to establish the platform’s diagnostic sensitivity, specificity, and clinical utility in real-world settings.

Second, while the platform is inherently compatible with multiplexed EV capture and biomarker detection, the simultaneous analysis of multiple EV subpopulations and biomarkers has not yet been systematically assessed. Before broader clinical implementation, a comprehensive evaluation of multiplexing performance, including potential assay interference and analytical robustness, will be required.

Future studies should focus on validating the platform in patient-derived samples across different disease contexts and on integrating the workflow with high-dimensional flow cytometry approaches to enable more comprehensive EV phenotyping and biomarker discovery.

## Authors contribution

Conceptualization, H.S.I., M.C.M, M.M., O.G.H., M.H., A.G., D.H., O.P.B.W.; Methodology, H.S.I., M.M., M.C.M., S.R., O.G.H., D.R.M., A.G., D.H., O.P.B.W.; Investigation, H.S.I., M.M., M.C.M., S.R., O.G.H., D.R.M.; Resources, D.H., O.P.B.W.; Writing – Original Draft, H.S.I.; Writing – Review and Editing, all the authors; Supervision, A.G., D.H., O.P.B.W.; Funding Acquisition, D.H., O.P.B.W.

## Acknowledgement

We would also like to acknowledge that CoPilot was used solely for proofreading and language editing purposes.

